# The Cell Death Census 2024

**DOI:** 10.1101/2024.04.04.588198

**Authors:** Mariam Miari, Elsa Regnell, Sonja Aits

## Abstract

Cell death plays a pivotal role in many physiological processes, such as cell homeostasis, embryonic development, immune defence and in the pathophysiology of numerous diseases, such as cancer, infections and degenerative diseases. However, the lack of a comprehensive and up-to-date resource on cell death regulators poses a significant challenge to researchers in the field. Existing databases are often limited in scope, differ in content and are updated irregularly. This deficiency impedes progress in understanding the intricate molecular mechanisms governing cell death and hampers the development of targeted therapies. To address this, we have performed a census of the existing cell death databases as well as the cell death-associated entries in the UniProt and Gene Ontology databases. To ensure high quality, we have focused on manually curated entries rather than those created from automatic prediction tools. The results have been consolidated into a joint database of the known cell death regulators, including both proteins and non-coding RNAs. The Cell Death Census 2024 results and the associated python code for database parsing, cleaning and merging is publicly available at https://github.com/Aitslab/CellDeathCensus/.

## Introduction

Cell death exists in accidental or regulated form (regulated cell death, RCD). Many forms of RCD, e.g. apoptosis, necroptosis, autophagic cell death, ferroptosis, have been described in the literature. In the past, these were thought of as distinct types of cell death but their tight interconnection has become increasingly obvious ^1, 2, 3, 4, 5^. Thus RCD is better thought of as a single process, executed through a network of redundant pathways and feedback loops.

Regulated cell death (RCD) in a physiological setting, often referred to as programmed cell death (PCD), is crucial for many of the key processes of life such as tissue and colonial homeostasis, tissue remodeling, embryonal development and immune defense ^6^. Consequently, both a lack of cell death where it is needed, or its aberrant activation contribute to a vast array of diseases in humans and other organisms. This makes cell death an important therapeutic target ^7, 8, 9^. Alterations in cell death can be caused by numerous genetic or epigenetic changes, toxic substances, pathogens or other cell stressors.

Despite the importance of RCD, the exact molecular pathways regulating and executing it in all of these situations is typically not yet elucidated in full. Consequently, while many therapeutic efforts aim to modulate RCD, it is not yet possible to do so by specifically targeting the cell death machinery. One exception is venetoclax, which inhibits the BCL2 apoptosis regulator (BCL2) inhibitor and is used in leukemia treatment ^10, 11^, providing proof of concept for the usefulness of such targeted drugs.

To capture the current state of knowledge on RCD, a number of manually curated databases of cell death regulators have been created ^12, 13, 14, 15, 16, 17, 18, 19, 20, 21^. In line with the traditional views on distinct cell death types, these are unfortunately typically limited in scope and most focus on apoptosis regulators. Even those with a broader scope, such as UniProt ^22^ and Gene Ontology ^23^ differ substantially in the listed cell death regulators. Furthermore, most of the databases are not updated regularly. To clear up this jungle of resources, we have performed a census of the existing databases and created a joint list over the known cell death-regulating proteins and non-coding RNAs. A similar effort has previously been made by the creators of the iPCD database ^15^, but unfortunately it has not been updated in several years and also left out some relevant databases. Our up-to-date census provides a clear overview over the state of knowledge in 2024, thereby facilitating further research on RCD.

## Methods

### Census procedure

Cell death databases were identified by Google search (search term “cell death database”) and from link lists in the websites of cell death-related organizations and databases. In addition, the GeneOntology and the UniProt database, which are widely used to look up functional associations with biological processes were included. To extract database content, bulk download files were used where they had been made available. Otherwise, database content was parsed from the database web site. Database entries not unambiguously related to cell death (e.g. autophagy regulators) or generated through automatic predictions (e.g. orthologue search) were excluded where possible. All data was downloaded on 2024-03-28 (ncRDeathDB V2 and FerrDb V2) or 2024-04-04 (all others).

Content from different databases included identifiers, symbols, primary and alternate names, functional information, species information and other descriptive data. Column headers and species names were reviewed manually and unified where possible to enable database merging (see Supplemental file 1 for column mapping). Names and symbols were pooled into a single synonym list as there was no consistent distinction between them in the databases. In addition, missing data that could be unambiguously identified from information on the database website, publication or other columns (e.g. species-related information, information about the cell death pathway and identifiers included in listed hyperlinks) was filled in.

For duplicate identification within individual databases, UniProt accession numbers (column ‘UniProt_AC’) were used where available. Alternatively, duplicates were assessed using gene symbols together with species information (columns ‘Symbol’ + ‘Species’ or ‘NCBI_TaxID’).

To ensure reproducibility and transparency, data extraction, cleaning and merging was performed using python code in a Colab notebook (Supplemental file 2). For the few exceptions where manual processing was needed, e.g. to download files, this is described in the notebook as well.

### Included databases

#### ApoCanD: Database of Human Apoptotic Proteins in the context of cancer ^12^

http://crdd.osdd.net/raghava/apocand/

This database contains 82 cancer-related genes involved in apoptosis.

#### DeathBase ^13^

http://www.deathbase.org/

This database contains genes involved in different cell death processes. As the bulk files available for download were incomplete, database content was parsed from the “Proteins” overview web page (http://www.deathbase.org/proteins.php, accessed 2024-03-21) and the “Full view” page (http://www.deathbase.org/edit_view.php, accessed 2024-03-21). Only the manually curated part, which contains proteins from human, fly, mouse, worm and zebrafish, was extracted, whereas the data for other organisms, resulting from homology searches, was excluded. Entries for which the Pathway was only “IMMUNITY” were also excluded.

#### FerrDB V2 ^14^

http://www.zhounan.org/ferrdb/current/

This database contains genes and compounds involved in ferroptosis. Database files for Driver, Suppressor, Marker, Unclassified sections were downloaded manually (http://www.zhounan.org/ferrdb/current/operations/download.html, accessed 20240325). The extended gene data from two more studies not yet included in the database ^24, 25^ was also downloaded from the FerrDB V2 website.

#### GO: Gene Ontology database ^23^

https://geneontology.org/

This general database contains information on genes related to all kinds of biological processes. Cell death-related GO terms were identified by manually reviewing the term tree on AmiGO 2, the official web tool for accessing the database (https://amigo.geneontology.org/) (Table 1). The selected high-level terms included the subterms linked to the UniProt keywords used when extracting UniProt data (Table 2). To extract all genes associated with the GO terms from AmiGO 2 a download link was generated after performing a manual search for the first term and then selecting fields of interest (Figure 1). Links for the other GO terms were created by id substitution in the link.

**Table 1.**
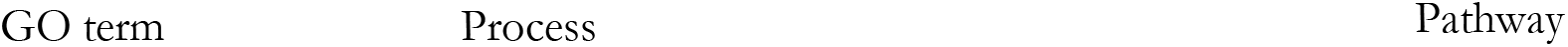
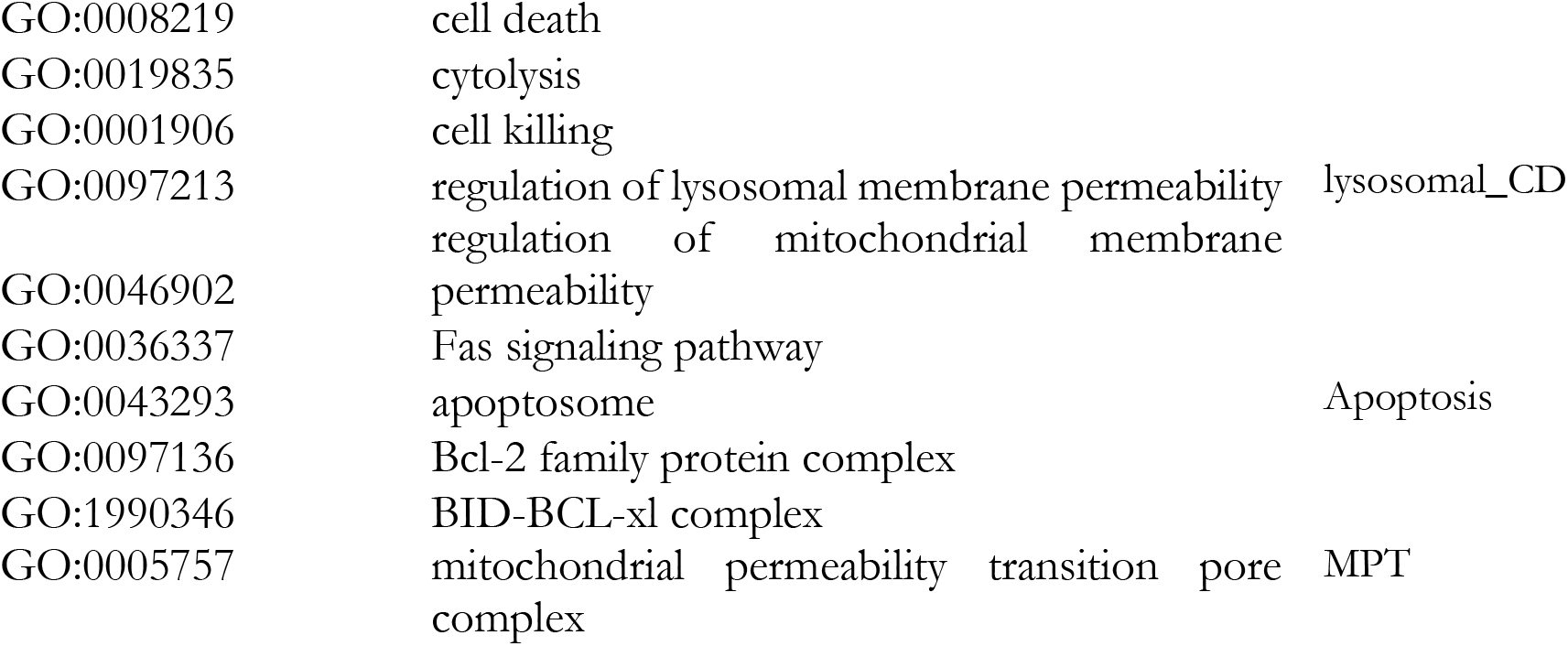
GO terms used in the census.

**Table 2.**
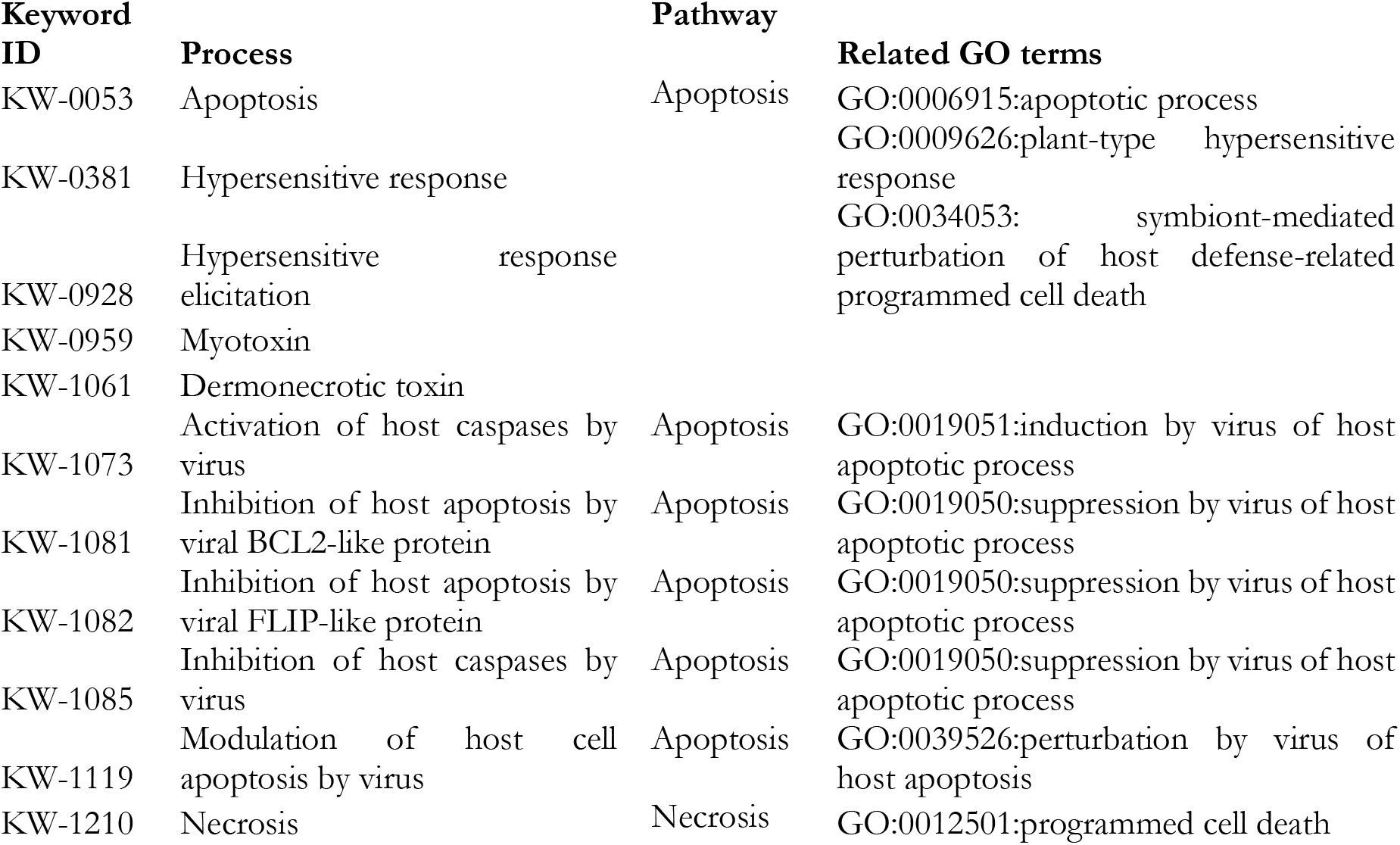
UniProt keywords used in the census.

**Figure 1.**
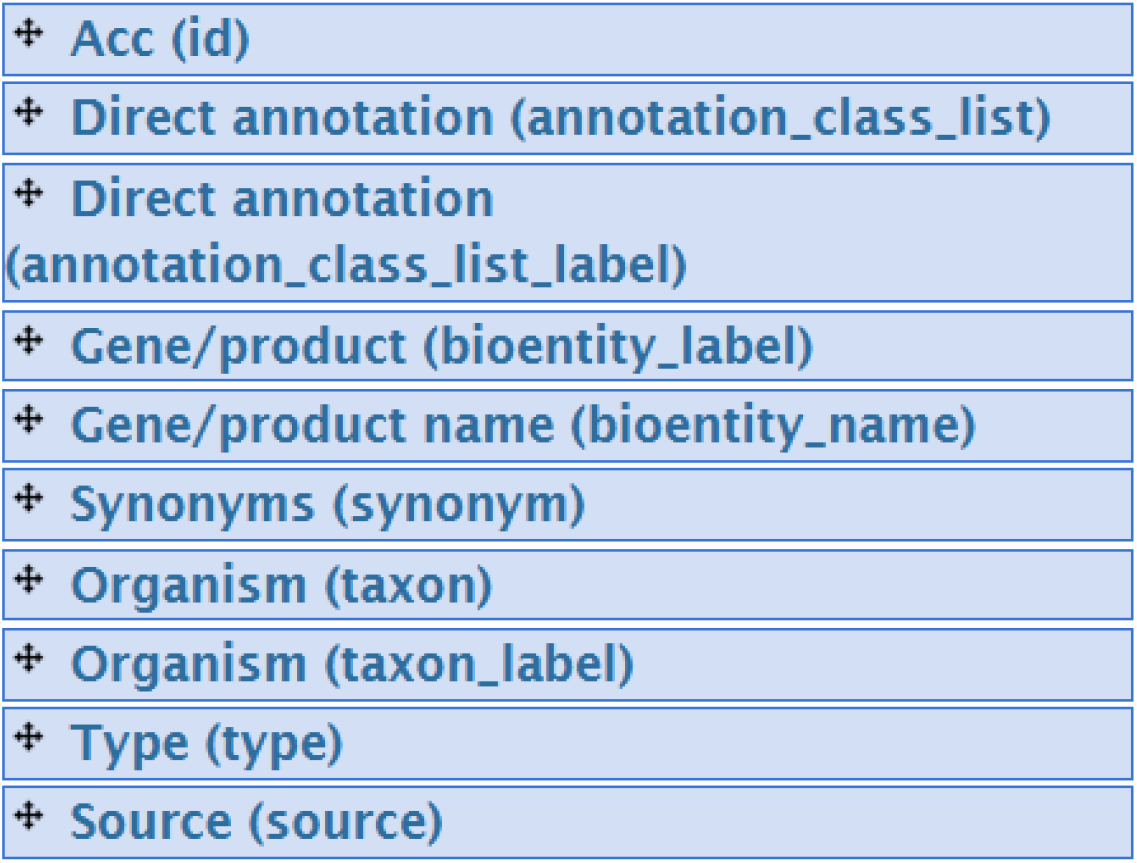
Fields extracted from the GO database.

#### iPCD: Integrated Annotations for Programmed Cell Death ^15^

http://ipcd.biocuckoo.cn/

This database contains genes involved in a very large variety of cell death types. Only the “Reviewed” part was extracted whereas the data generated by orthologue search (“Unreviewed”) was excluded. Entries only related to Autophagy were also excluded.

#### lncPCD ^16^

*http://spare4.hospital.studio:9000/lncPCD/*

This database contains lncRNAs associated with apoptosis, autophagy, ferroptosis, necroptosis and pyroptosis and their disease associations. Entries only related to Autophagy were excluded.

#### MCDB: Mitotic Catastrophe Database ^17^

http://www.combio-lezhang.online/MCDB/index_html/

This database contains genes and compounds related to mitotic catastrophe.

#### ncRDeathDB, Release 2.0 ^18^

http://www.rna-society.org/ncrdeathdb/

This database contains miRNAs, lncRNAs and snoRNAs related to apoptosis, necrosis and autophagy and their target genes. Both non-coding RNAs and target genes were extracted. Autophagy-related entries were excluded.

#### RCDMap: Comprehensive Map of the Regulated Cell Death Signaling Network ^19^

https://navicell.vincent-noel.fr/pages/maps_rcd.html

As the data was no longer accessible on the main website, the version deposited in Minerva was downloaded by manual export (https://acsn-curie.lcsb.uni.lu/minerva/index.xhtml?id=Regulated_Cell_Death, accessed 20240404).

#### UniProt, Release 2024-02 ^22^

https://www.uniprot.org/

This general database contains information on proteins related to all kinds of biological processes. All UniProt keywords were downloaded and manually reviewed to identify those linked to cell death. For each keyword all “Reviewed” entries, corresponding to the SwissProt part of UniProt, were extracted. Species names were typically in the format Latin name (Common name). For the most frequent species, common names were removed to enable merging with other databases.

#### yApoptosis ^20^

http://www.ycelldeath.com/yapoptosis/

This database contains genes related to apoptosis in yeast.

#### XDeathDB ^21^

https://pcm2019.shinyapps.io/XDeathDB/

This database contains human genes related to 12 modes of cell death and information on associated diseases and drugs. After searching in the Cell Death Engine section for cell death modes “All” and disease types “All” the csv file with all entries was downloaded manually. Entries displaying “Autophagy”, “Proliferation” or no value as cell death mode were excluded.

### Excluded databases

#### Apoptosis DB ^26^

http://www.apoptosis-db.org

This database contained proteins related to apoptosis. It is no longer publicly accessible.

#### BCL2DB: BCL2 DataBase ^27^

https://bcl2db.lyon.inserm.fr/

This database was not available for bulk download and undergoing a major revision at the time of the census (personal communication with the developer Christophe Combet, 20240321).

#### CASBAH : The CAspase Substrate dataBAse Homepage ^28^

https://bioinf.gen.tcd.ie/casbah/

This database contained caspases and their substrates. It is no longer publicly accessible.

#### CASBASE: The Database for Caspases & Substrates

http://origin.bic.nus.edu.sg/casbase/

This database contained caspases and their substrates. It is no longer publicly accessible.

#### Cell Death Proteomics (CDP) database ^29^

http://celldeathproteomics.uio.no/

This database contained proteins related to cell death. It is no longer publicly accessible.

## Results

### Cell death-related databases greatly vary in scope, size and update frequency

Seven cell death-specific and two general gene/protein annotation databases were included in the census (Table 3). The Bcl2DB database was excluded as it is currently undergoing a major revision and lacks bulk download options. This will hopefully be completed in time for the next census update. In addition, four databases reported in the literature were found to be no longer accessible: Apoptosis DB, CASBAH, CASBASE and Cell Death Proteomics database.

**Table 3.**
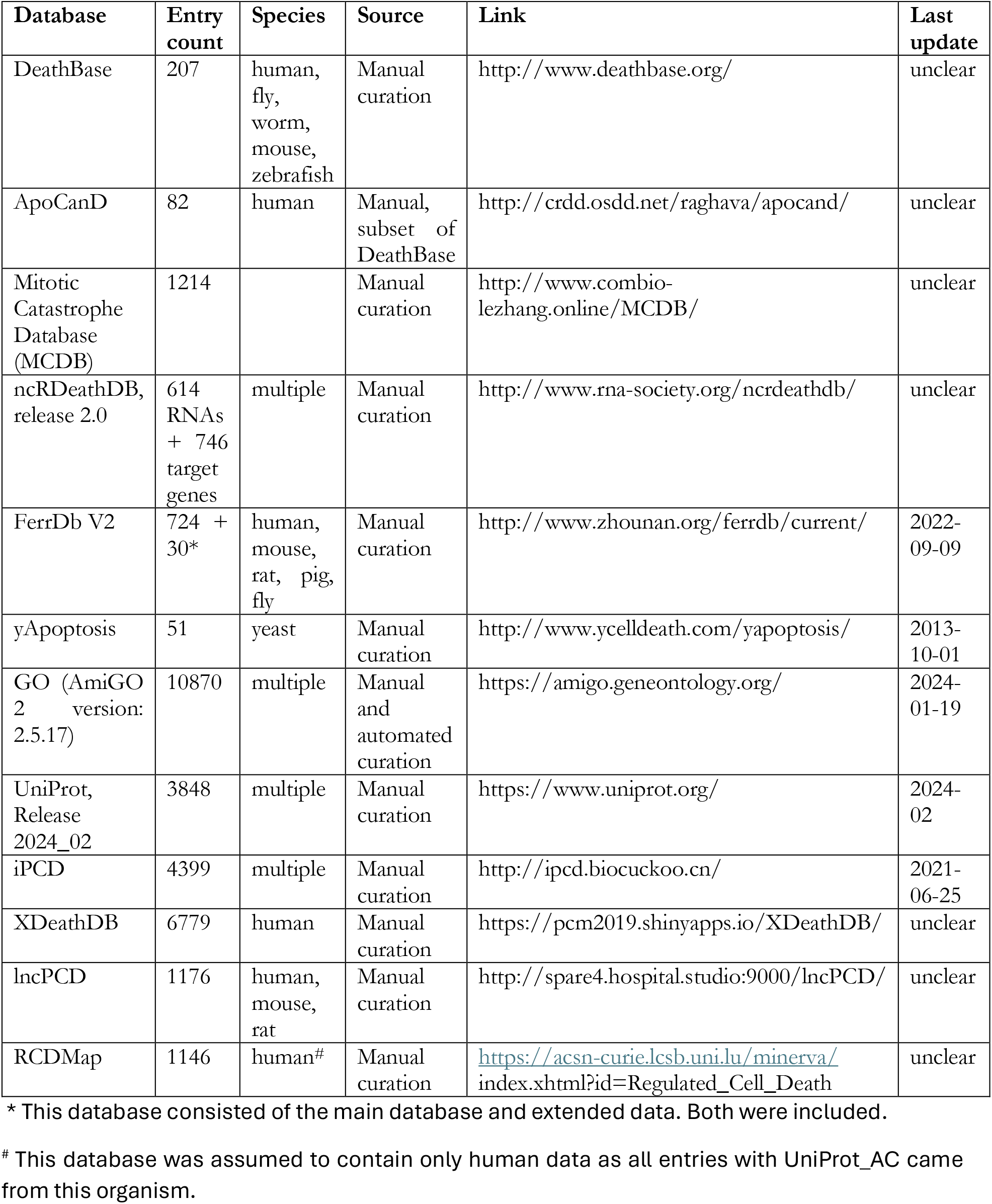
Overview over the databases in the Cell Death Census 2024.

When examining the content of the available cell death-related databases, large variations were observed regarding the types of cell death, species, size and update frequency. While the large general databases UniProt and GO were not limited to a specific cell death type, only one of the specialized cell death databases, iPCD had a similarly broad scope. A similar limitation in scope was seen on the species level. Both the iPCD and the DeathBase also have a section with many additional species and entries, generated by orthologue prediction but as we wanted to focus on high quality manually reviewed data this data was not included. The number of entries in each database, after removal of duplicates ranged from 51 (yApoptosis) to 10870 (GO). However, a large part of the GO database is not manually curated. As it was not immediately obvious which entries in GO stemmed from automated annotation, these were not excluded for this database.

Of the databases which reported their last update on the website, which several did not, only UniProt and GO were updated regularly. iPCD, the only other database comparable to our efforts, was last updated in 2021, highlighting the need for this new census.

### The Cell Death Census 2024 gathers a large number of cell death regulators but species coverage is very uneven

To merge the data obtained from the different databases, column names were manually reviewed and unified where suitable. Missing data was filled in where unambiguously possible. For example, we added species information listed on the website and extracted identifiers contained in hyperlinks. In total, we obtained 49544 records from the census (Supplemental file 1). This included a large number of partially duplicate records. As the naming was inconsistent between databases it was impossible to fully merge the duplicates without extensive manual review. It was therefore not attempted.

Nevertheless, we tried to obtain an estimate of the unique number of regulators in the census dataset. For this purpose, we first examined the presence of commonly used identifiers (Table 4) that could be used for merging. Despite the ambiguity, symbols (column ‘Symbol’) were the most frequent identifiers, followed by UniProt accession number (column ‘UniProt_AC’), which covers only proteins. When merging the Census results on UniProt_AC, we obtained 14746 unique records (Supplemental file 4).

**Table 4.**
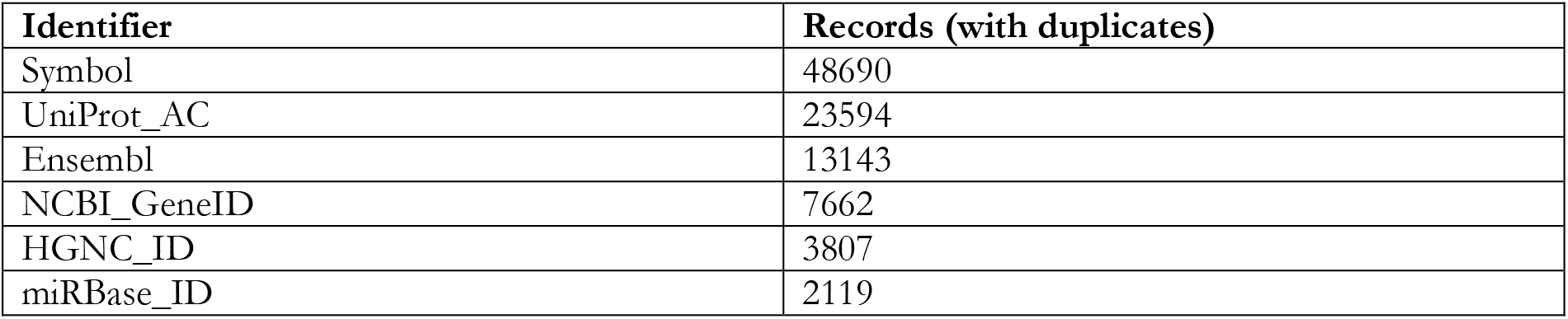
Frequency of common identifiers in the Cell Death Census (45012 records)

An alternative strategy to merge duplicates is to merge on symbol and species information combined, thereby reducing ambiguity. For this, we first filled in missing values for Species and NCBI_TaxID based on the existing pairs in the dataset. When counting the occurrence of each species, it was clear that most records represent human data (Table 5). After merging on the combined Symbol and Species, we obtained 25555 unique records (Supplemental file 5). Similar results were seen after merging on Symbol and NCBI_TaxID which resulted in 25329 unique records (Supplemental file 6.

**Table 5.** Frequency of the most common species terms in the Cell Death Census.

| Species | Count |
| --- | --- |
| homo sapiens | 25594 |
| mus musculus | 5485 |
| rattus norvegicus | 2426 |
| bos taurus | 932 |
| canis lupus familiaris | 882 |
| sus scrofa | 839 |
| danio rerio | 596 |
| strongylocentrotus purpuratus | 558 |
| gallus gallus | 542 |
| drosophila melanogaster | 471 |
| xenopus laevis | 330 |
| caenorhabditis elegans | 319 |
| xenopus tropicalis | 318 |
| oryzias latipes | 258 |
| arabidopsis thaliana | 237 |
| macaca mulatta | 212 |
| pan troglodytes | 207 |
| equus caballus | 205 |
| felis catus | 204 |
| gorilla gorilla gorilla | 203 |
| branchiostoma floridae | 183 |
| lepisosteus oculatus | 180 |
| daphnia pulex | 178 |
| monodelphis domestica | 157 |
| ornithorhynchus anatinus | 142 |
| triticum aestivum | 140 |
| saccharomyces cerevisiae s288c | 132 |
| pongo abelii | 128 |
| saccharomyces cerevisiae | 125 |
| anolis carolinensis | 115 |
| candida albicans sc5314 | 112 |
| nakaseomyces glabratus cbs 138 | 109 |
| candida dubliniensis cd36 | 102 |
| candida parapsilosis cdc317 | 102 |
| anopheles gambiae | 97 |
| nematostella vectensis | 95 |
| helobdella robusta | 95 |
| [candida] auris | 91 |
| caenorhabditis briggsae | 90 |
| tribolium castaneum | 86 |
| arabidopsis thaliana (mouse-ear<br>cress) | 82 |
| brassica napus | 77 |
| pristionchus pacificus | 73 |
| dictyostelium discoideum | 66 |
| nicotiana tabacum | 61 |
| ciona intestinalis | 60 |
| gossypium hirsutum | 58 |
| capsicum annuum | 55 |
| glycine max | 52 |
| ixodes scapularis | 49 |

**Table 6.**
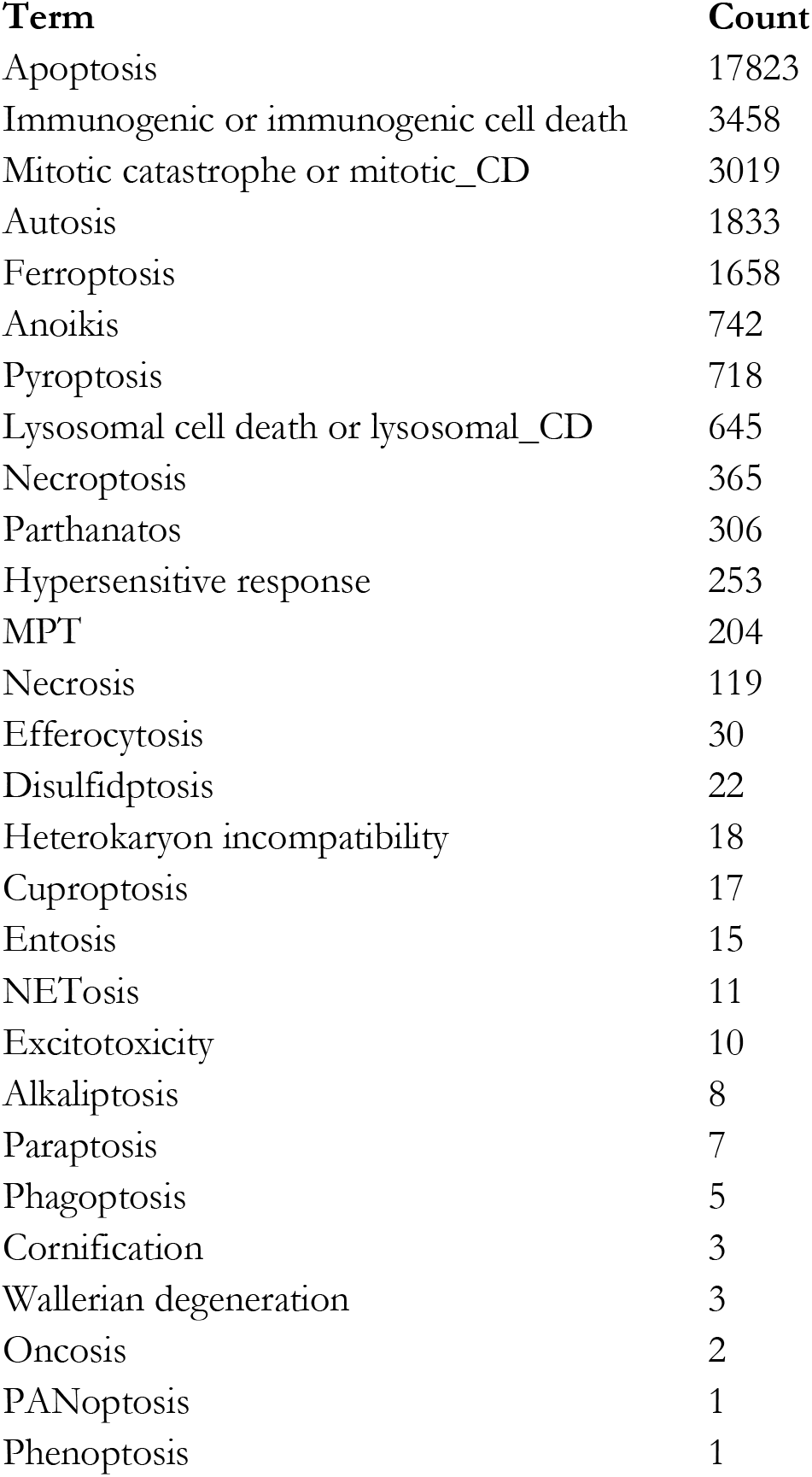
Frequency of cell death terms in the Cell Death Census.

### Existing cell death databases are strongly biased towards apoptosis

We next examined the frequency of different cell death types. A strong bias towards apoptosis was observed. One notably absent term was “autophagic cell death”. Several databases had included autophagy regulators, but these were excluded as the databases did not distinguish between autophagic cell death regulators and pro-survival autophagy regulators.

## Discussion

It is important for researchers to be aware of the state of the current knowledge in their field. Therefore, many bioinformatics databases have been produced to list genes, proteins and/or non-coding RNAs linked to specific processes. These databases can be populated by manual curation or algorithms. Such databases are for example used when estimating changes in specific pathways in omics experiments or when trying to piece together signaling pathways. Within the field of cell death research, several such databases have been compiled with manually curated data. These vary greatly in species, cell death processes, gene types, number of entries and other aspects. The scatteredness of this information makes it difficult for researchers to make use of it in full. There has previously been an effort by the iPCD database to combine some of this data but unfortunately general databases that are widely used in pathway analysis were left out. The database has also not been updated since 2021. To unite the existing information on cell death regulators we therefore conducted a census of the available databases, also including GO and UniProt, two of the most widely used general databases with functional annotations. All results as well as two Colab notebooks (Supplemental file 1-3) to repeat the analysis are made publicly available. The latter enables cell death researchers to repeat the census in an almost fully automated manner. Frequent updates of the census do not seem to be necessary, however, as most of the surveyed databases had not been updated in years.

The Census revealed a large number of reported cell death regulators. However, there was a strong bias towards human apoptosis-related protein-coding genes. Future curation efforts should therefore aim to include genes from a broader range of species and alterative cell death types as well as non-coding RNAs.

One of the largest problems encountered in the data analysis was the inconsistent use of unique identifiers. Indeed, many entries were only listed with the highly ambiguous gene symbol. This makes it impossible to fully merge the content of different databases without extensive manual work. We strongly recommend that database designers list several common identifiers for each entry, e.g. UniProt_AC, HGNC_ID and NCBI_GeneID. Likewise, species should be unambiguously identified with the widely used NCBI_TaxID.

Another problem was that several databases were no longer accessible or lacked bulk download options. Database developers should consider uploading a copy of their database and all updates to a general research data repository. This would ensure that manual curation efforts are not lost when database developers can no longer maintain the online presence of their database. At the same time, deposited datasets will also receive a DOI and version number, making it easier for other researchers to cite the data in a clear manner.

A full list of recommendations for database developers based on our experience with the Census can be found in Box 1.

### Box 1.

**Recommendations for database development and maintenance**

1. At the time of release and with every update, a copy of the database should be deposited in a general research repository (e.g. zenodo) where it will receive a DOI and version number. This DOI should be included in the research article describing the database.
2. A clear description of the database scope (e.g. gene type, species, cell death type), curation procedure, manual annotation guidelines, external sources, files and column headings should be published on the database page and deposited in a general research repository together with the database content. For automated content collection, e.g. by orthologue search or import from other databases, a clear description of the procedure and all settings, and/or the code used in the process, should be provided.
3. The date of the last update and database version should be highlighted on the landing page of the database.
4. An option for automated bulk download of the database content should be provided.
5. Genes should be identified unambiguously with several general identifiers, e.g. UniProt_AC + HGNC_ID + NCBI_GeneID. Ensure that the identifiers are still active at the time of database release and update with each release.
6. Species should be identified unambiguously with NCBI_TaxID + Linnean name.
7. PMIDs should be provided so end users can review the evidence used during manual curation.
8. Delimiters used to separate content within a column and between columns should be different and used consistently.
9. Empty values should be indicated clearly and consistently.

## Acknowledgement

This study was supported by the Swedish Research Council, the Wallenberg AI, Autonomous Systems and Software Program – Humanities and Society (WASP-HS) and Data-driven life science (DDLS) program, the Swedish Research Council for Sustainable Development (FORMAS), and the Lund University Sustainability Fund.

We also acknowledge the following research environments and networks which support our work: AI Lund, AIR Lund, LTH Profile Area: AI and Digitalization, LTH Profile Area: Engineering Health, EpiHealth: Epidemiology for Health and PhenoTarget.

We thank Rafsan Ahmed for critical reading of the text.

## CRediT author statement

Mariam Miari: Data Curation, Software

Elsa Regnell: Data Curation

Sonja Aits: Conceptualization, Methodology, Software, Validation, Formal analysis, Investigation, Resources, Data Curation, Writing - Original Draft, Writing - Review & Editing, Visualization, Supervision, Project administration, Funding acquisition

## Supplemental data

Supplemental files are available on https://github.com/Aitslab/CellDeathCensus/SupplementalData

Supplemental file 1. Cell Death Census 2024 – zip file with all merged data

Supplemental file 2. Cell Death Census 2024 – Data collection colab notebook

Supplemental file 3. Cell Death Census 2024 – Data analysis colab notebook

Supplemental file 4. Cell Death Census 2024 – unique entries merged on UniProt_AC

Supplemental file 5. Cell Death Census 2024 – unique entries merged on Symbol + Species

Supplemental file 5. Cell Death Census 2024 – unique entries merged on Symbol + NCBI_TaxID

